# Multisite generalizability of schizophrenia diagnosis classification based on functional brain connectivity

**DOI:** 10.1101/141192

**Authors:** Pierre Orban, Christian Dansereau, Laurence Desbois, Violaine Mongeau-Pérusse, Charles-Édouard Giguère, Hien Nguyen, Adrianna Mendrek, Emmanuel Sti, Pierre Bellec

## Abstract

Our objective was to assess the generalizability, across sites and cognitive contexts, of schizophrenia classification based on functional brain connectivity. We tested different training-test scenarios combining fMRI data from 191 schizophrenia patients and 191 matched healthy controls obtained at 6 scanning sites and under different task conditions. Diagnosis classification accuracy generalized well to a novel site and cognitive context provided data from multiple sites were used for classifier training. By contrast, lower classification accuracy was achieved when data from a single distinct site was used for training. These findings indicate that it is beneficial to use multisite data to train fMRI-based classifiers intended for large-scale use in the clinical realm.

## 1. Introduction

Psychiatrists and other mental health professionals could benefit in the not-so-far future from neuroimaging-based classification tools to assist diagnosis and prognosis in mental illness (Huys et al., 2016). Recent developments in the neuroimaging field have led to a shift from group comparisons based on averaging across subjects to machine learning techniques making prediction at the individual level (Dubois and Adolphs, 2016). In this approach, the emphasis is put on the ability of an algorithm to classify individuals into clinical categories with good generalizability to unseen subjects. Over the last decade, hundreds of studies have successfully classified various psychiatric and neurological disorders based on in vivo brain imaging (reviewed in Arbabshirani et al., 2016; Wolfers et al., 2015). For instance, Arbabshirani et al. (2016) identified 30 published studies that distinguished schizophrenia patients from healthy controls with an average accuracy of 83% using functional magnetic resonance imaging (fMRI), either under task or rest states.

To date, however, the vast majority of classification works in mental illness were performed in a research context, using data from single sites of acquisition. Such findings may not generalize to large-scale clinical settings, with patients being scanned at widely-spread sites and possibly under various mental states. In most cases, the performance of classifiers was only assessed for unseen, test subjects with the exact same characteristics as the sample used for training. Yet, using gender as a proof-of-concept target variable, there was initial evidence that classifiers only poorly generalize to data drawn from other site samples (Huf et al., 2014). The inclusion of data from multiple sites during training improved the classifier performance for data of unseen sites.

In schizophrenia, a study pooling fMRI data from two distinct scanning sites reported similar prediction accuracy levels irrespective of whether test data were drawn from the dataset used for training or not, thus suggesting good generalizability (Skaåtun et al., 2017). However, this result appears at odds with a recent fMRI study in autism that showed poorer accuracy for inter-site than intra-site training/test configurations, depending on the ratio of training set used (Abraham et al. 2017). In the case of inter-site testing, data pooled from 4 sites were used for training the classifier, which was tested on data from a fifth site. Yet, none of these two studies specifically evaluated whether using multisite training data could compensate to some extent for the deleterious effect of inter-site testing, by assuming the actual presence of such an effect. In the present work, we sought to address this question based on fMRI brain connectivity in schizophrenia. Since it is impossible to completely control the variations in mental states in realistic clinical situations, we further promoted the complexity of the classification problem by including data obtained in distinct cognitive task conditions across sites. Mass univariate findings have indicated that cognitive state does not further impact on the nature of functional brain connectivity alterations in schizophrenia (Kaufmann et al., 2016; Orban et al., 2016). However, the potential influence of cognitive context on classification performance in a multivariate analysis should not be rejected.

## 2. Methods

### 2.1 Datasets

Brain imaging data from 6 independent studies were obtained through either the SchizConnect and OpenfMRI data sharing platforms (http://schizconnect.org; https://openfmri.org) or local scanning (Çetin et al., 2014; Gollub et al., 2013; Kogan et al., 2016; Orban et al., 2016; Poldrack et al., 2016; Wang et al., 2016). The 6 datasets differed in terms of both scanning site and cognitive context during fMRI data acquisition (resting-state, emotional memory, Sternberg item recognition paradigm, N-back, task-switching and oddball tasks). Classification analyses included fMRI data from 382 subjects, 191 patients diagnosed with schizophrenia and 191 healthy controls. Subjects provided informed consent to participate in their respective studies and ethics approval was obtained at the site of secondary analysis (Centre de Recherche de l’Institut Universitaire de Gériatrie de Montréal, Montréal, Canada).

### 2.2 Subjects matching

Sample size differed between sites (N = 84, 82, 70, 62, 50 and 34). Site samples were obtained after subjects were selected in order to ensure even proportions of schizophrenia patients and controls within each site (N = 42, 41, 35, 31, 25 and 17 subjects per group) and to reduce between-group differences with regards to gender ratio (75% vs 73% males in controls vs. schizophrenia patients), age distribution (32.3 ± 9.8 vs. 33.4 ± 9.5 years old) and motion levels (average frame displacement = 0.15 ± 0.05 vs 0.17 ± 0.06, see Data preprocessing). Matching of schizophrenia and control subjects was achieved based on propensity scores, using the Optmatch R library version 0.9–7 (https://cran.r-project.org/web/packages/optmatch/index.html). The propensity score associated with each participant was defined by the conditional probability of being in the clinical or control group given the confounding covariates (gender, age and motion). Propensity scores were then used to balance those covariates in the two groups. Although we took great care in matching participants with respect to these factors of no interest, it is very likely that other confounds such as medication in schizophrenia patients impacted the reported findings.

### 2.3 Data preprocessing

Brain imaging data preprocessing and extraction of functional brain connectomes were performed with the NeuroImaging Analysis Kit version 0.12.17 (NIAK, http://niak.simexp-lab.org). Briefly, preprocessing included slice timing correction, estimation of rigid-body motion within the functional runs, nonlinear coregistration of the structural scan in stereotaxic space, individual coregistration between structural and functional scans, resampling of the functional scans at 3mm isotropic resolution in stereotaxic space, scrubbing of volumes with excessive motion (frame displacement greater > 0.5 mm), regression of confounds (slow time drifts, average of conservative white matter and cerebrospinal fluid masks and motion parameters), and smoothing of functional volumes with a 6 mm isotropic Gaussian blurring kernel. A detailed description of the preprocessing pipeline can be found at http://niak.simexp-lab.org/pipe_preprocessing.html.

Individual functional connectomes included 2016 functional connections between 64 brain parcels. The functional brain parcellation was previously obtained by conducting a bootstrap analysis of stable clusters (BASC, Bellec et al. 2010) on an independent fMRI dataset of 200 healthy young subjects (https://doi.org/10.6084/m9.figshare.1285615.v1). In each schizophrenia or control participant, the time series of a brain parcel consisted in the average of the voxel signals in the parcel. Connectivity measures between pairs of parcels were defined by Pearson product-moment correlation coefficients. Individual connectomes were parcel by parcel (64 x 64) symmetrical matrices that summarized connectivity levels in the whole brain. Lower triangular matrices were then vectorized for all subjects in order to form a subject by connections (382 x 2016) matrix.

### 2.4 Data analysis

Classification analyses were performed with a linear support vector machine (SVM) algorithm, as implemented in the SciKit-Learn python library version 0.18.1 (Abraham et al., 2014). The SVM classifier, a supervised classification algorithm, represented subjects as points in space, mapped so that the subjects of the separate clinical labels were divided by a clear gap (called a margin) that was as wide as possible. The hyperparameter C of the SVM was optimized using nested cross-validation. Each model used the residuals from a regression of confounding variables (gender, age and motion parameters) across connections estimated from the subjects selected for training the model. The evaluation metrics were computed using four main values, namely the number of true and false positive (TP, FP) as well as true and false negatives (TN, FN). Sensitivity was defined as TP/(TP+FN), specificity as TN/(TN+FP) and accuracy as (TP+TN)/(TP+FP+TP+FN). The main analyses evaluated the impact on classification accuracy of the number of site(s) (1, 2, 3, 4 or 5) included in the training set. We evaluated this impact in situations where the test set included only subjects from the same site(s) used during training (intra-site test with 10-fold cross validation) or, alternatively, situations where the test set included only subjects from sites not used during training (inter-site test with “leave-site-out” cross validation). Cross validation ensured that the subjects used for training were never used in the test phase.

The statistical significance of changes in accuracy levels as a function of the number of sites used for training and whether data used for testing were drawn from the same dataset(s) used for training (intra-site vs inter-site) was assessed with binary logistic regressions using the GLM function in R version 3.2.5. These analyses relied on the prediction of categorical outcomes (hit/miss data) based on predictor variables (number of sites used for training, intra-site vs inter-site). Significance threshold in the different contrasts was set at p < 0.05.

Complementary analyses were conducted. First, we explored differences in whole brain connectivity between schizophrenia patients and controls using mass univariate statistics for the various training site combinations. Similarly for multivariate classification analyses, we extracted feature weights separately for all site combinations. We then examined the level of correspondence across site combinations for both univariate and multivariate analyses. Second, we aimed at demonstrating the presence of multivariate site effects on functional brain connectivity. To this end, we determined accuracy levels for the classification of scanning sites by performing separate SVM analyses for all pairs of sites, using 10-fold cross validation as in the main analyses.

## 3. Results

### 3.1 Correspondence across site combinations

We first report patterns of functional brain dysconnectivity in schizophrenia patients based on mass univariate statistics. For the sake of interpretability, the 64 brain parcels were sorted in relation to 7 large-scale brain networks from the same multiscale functional brain atlas (Figure 1a,b). When pooling data from all subjects and sites, a connectome-wide association analysis revealed widespread decreased connectivity in schizophrenia patients (Figure 1c), with 769 out of 2016 connections exhibiting a significant effect after false discovery rate correction (q^FDR^ < 0.05). Differences between schizophrenia patients and controls were further examined separately for each unique combination of 1 to 5 training sites (61 possibilities: 1, 2, 3, 4, 5, 6, 1-2, 1-3, 1-4, …, 1-2-4-5-6, 1-3-4-5-6, 2-3-4-5-6). Results revealed high variability in the nature of mass univariate effects across training site combinations, with small correlation between them when there was no overlap between site combinations (Figure 1c). By contrast, large correlations were observed when site combinations overlapped. Weight matrices, which indicate for each connection the importance of that connection in the decision process, were also extracted for the whole sample as well as each site combination in multivariate classification analyses (Figure 1d). The correspondence between site combinations mimicked the patterns of correlations seen for univariate analyses, with a large correlation between weights for site combinations that overlapped but a small correlation otherwise.

**Figure 1.**
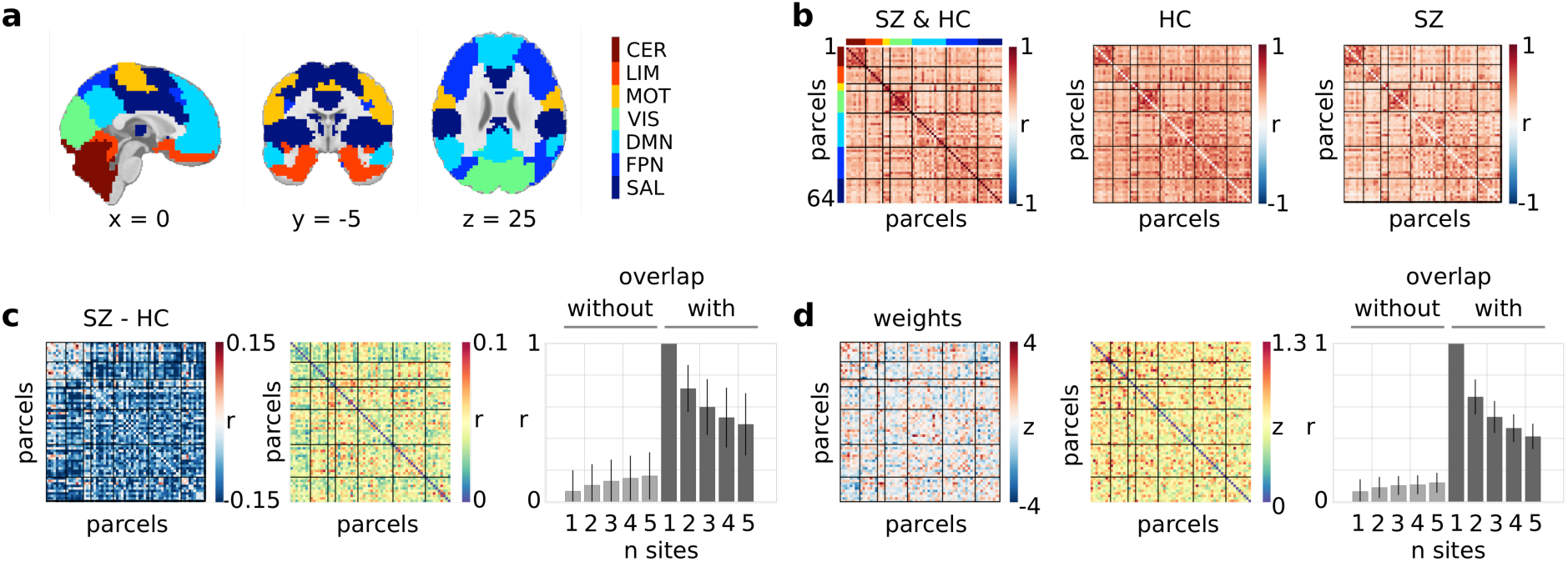
Correspondence between connectomes. Sorting the 64 brain parcels in relation to 7 large-scale brain networks (a) allowed to best reveal the structure of whole-brain connectivity in both schizophrenia patients and controls, here shown when combining data from all sites (b). Results for univariate analyses (c). The left panel reports mean differences in connectivity between schizophrenia patients and controls pooled across all sites. The middle panel shows variations (standard deviation) of between-groups differences across the various site combinations. The right panel reports correlations of univariates effects of schizophrenia between single sites of reference (sites 1 to 6) and various site combinations as a function of whether site combinations included the sites of reference and the number of sites included in the combination (mean and standard deviation across the 6 reference sites). Results for multivariate classification analyses are similarly organized (d). The left panel provides the normalized weights obtained when pooling all subjects from all sites. The middle panel indicates how these weights vary across site combinations (standard deviation). The right panel provides correlations of weight matrices between single sites of reference and various site combinations. Abbreviations for networks are as follows: CER, cerebellum; VIS, visual; LIM, limbic, MOT, motor; SAL, salience; FPN, fronto-parietal; DMN, default-mode.

### 3.2 Classification findings

Classification of sites was performed with high accuracy (84%), indicating a significant multivariate impact of scanning site on functional brain connectivity. Training on data from a single site led to a poor generalization of diagnosis classification to subjects drawn from another site, i.e. classification accuracy was much lower in the inter-site than intra-site configuration when only one site was used for training (p<0.005). However, increasing the heterogeneity of the training set by including data from different sites improved accuracy of the classifier applied to another unknown scanning site and cognitive context (p < 5x10^-8^) (Figure 2a). This compensatory effect was such that inter-site classification reached similar accuracy performance to intra-site classification when 5 different sites were used for classifier training (p = 0.56), thus suggesting excellent generalization in this context. The benefit of using heterogeneous training data when classifying subjects drawn from the same sites as the training set was much more moderate than for the inter-site training-test configuration, yet was significant (p <0.05). Formal testing of an interaction effect revealed a significant effect (p < 0.05), thus demonstrating that improved generalization on novel sites following multisite training was not merely a consequence of increasing sample size.

**Figure 2.**
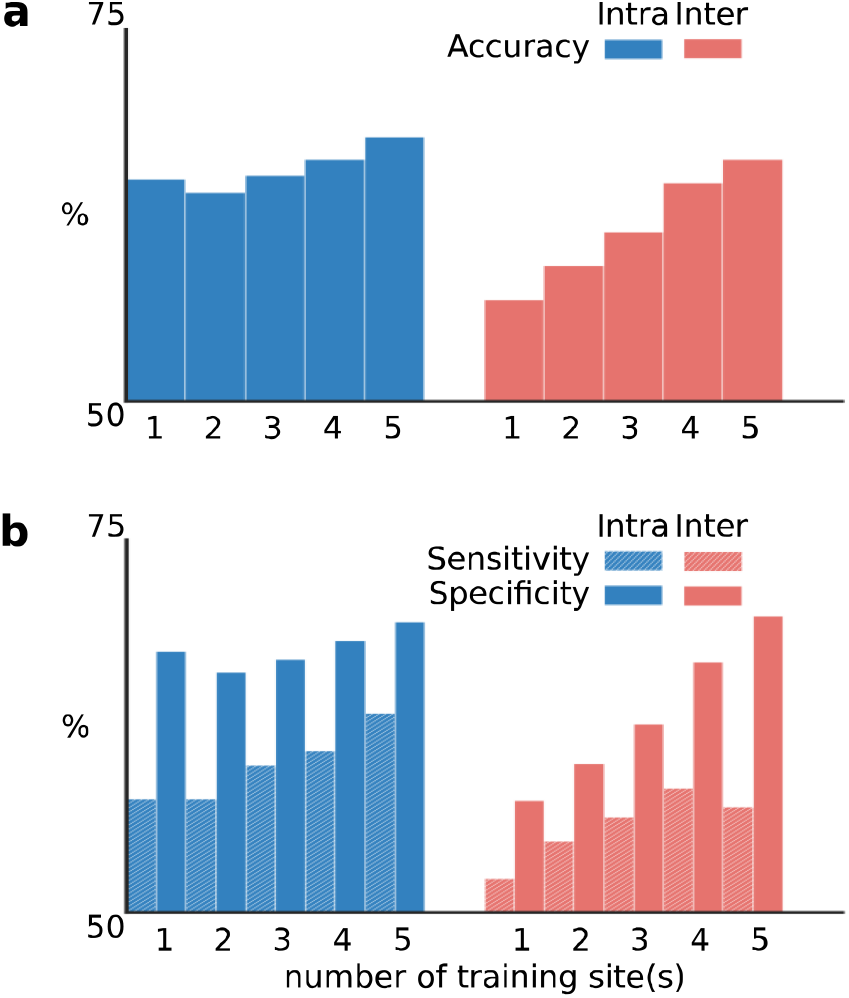
Classification findings. Histograms show the percentages of classification accuracy (a) as well as sensitivity and specificity (b) as a function of the number of sites (1, 2, 3, 4 or 5) from which individual connectomes were drawn for training, and whether testing was performed on subjects drawn from the same site(s) as during training (intra-site test) or not (inter-site test).

## 4. Discussion

The present findings highlight a prerequisite for an optimal translation of classification tools from the research to clinical realm. Namely, classifier training should be performed on data that are sufficiently representative of sites and/or mental state variations in order to generalize well for large-scale clinical use. In particular, the accuracy scores reported in most of the existing literature should be interpreted with caution, as they only reflect within-site generalizability and may therefore overestimate the accuracy.

Mass univariate analyses evidenced brain dysconnectivity across the entire brain, with significant effects in over a third of brain connections distributed in various large-scale brain networks, from cognitive to primary sensory networks. Abnormally decreased rather than increased functional connectivity in schizophrenia is largely consistent with previous reports in the literature (Pettersson-Yeo et al., 2011). With close to 200 schizophrenia patients and 200 controls, our fMRI connectome-wide association analysis is one of the largest to be reported to date. The present work and previous similar studies conducted in schizophrenia (Cheng et al., 2015; Schilbach et al., 2016; Skatun et al., 2017) underscore the utility of pooling data across multiple sites of acquisition in order to achieve higher sample size and more reliable findings. The marked variability in dysconnectivity patterns detected in each site separately could induce a deleterious effect of multisite data pooling on statistical power as compared to data obtained at a single site. However, there is support to claim that the initial deleterious effect of multisite data pooling can be mitigated by the increase in sample size and number of sites in the context of intra-site mass univariate as well as multivariate analysis (Dansereau et al., 2017). This is in line with our findings suggesting that a similar compensating effect can be obtained using multisite data aggregation in inter-site multivariate prediction, a configuration that is most likely to be found in clinical settings.

Multivariate classification analyses indicate that increasing sample size through multisite data pooling increased diagnosis prediction in schizophrenia, although this effect was of small amplitude. More critically, an additional benefit of including heterogeneous data was that the classifier generalized better to data that were not represented during training, neither in terms of scanning site nor mental content. This demonstration is concordant with a previous report that classified gender as a proof-of-concept application (Huf et al., 2014), and underscores the benefit of pooling multisite data for the purpose of generalizability and clinical use. The observed gain of almost 10% in classification accuracy is appreciable. It is nonetheless noteworthy that the highest accuracy of schizophrenia diagnosis classification was below 70%, which precludes the immediate translation of such machine learning tools in the clinical realm. Beyond the fact that most classification work has to date investigated within-site generalizability, it is notable that most studies relied on small samples. This is likely to be accompanied by a publication bias by which only the most significant findings were published. While the average classification of schizophrenia diagnosis over 30 published studies is above 80%, it was accordingly shown that studies with a large sample size in fact reported lower classification accuracy (Arbabshirani et al., 2016). Besides, low classification accuracy is very likely dependent on the ill-definition of clinical labels, as schizophrenia is a highly heterogeneous psychiatric disorder (Kapur et al., 2012). The stratification of patients into more homogeneous neurobiological subtypes, beyond clinical symptoms, will likely define more precise labels that will lead to improved classification of their diagnosis. The characterization of mental illness heterogeneity through the identification of such different biotypes, in particular based on fMRI brain connectivity, is a topic of burgeoning research in various psychiatric disorders (Costa Dias et al., 2015; Drysdale et al., 2016; Gates et al., 2015).

It is anticipated that neuroimaging-based classification will ultimately assist psychiatrists in not only diagnosis but also prognosis and theragnosis in mental illness, including schizophrenia. The future integration of classifiers into mental health care will require studies with dramatically increased sample size (Dubois and Adolphs, 2016). Most studies indeed suffer from insufficient data, possibly resulting in biased accuracy estimation, under-representation of mental illness heterogeneity and unstable findings (Arbabshirani et al., 2016). Future work will also need to develop novel algorithms with improved capabilities and to better define clinical labels. The present work identifies one specific parameter that will facilitate an optimal translation of supervised machine learning into clinical practice, namely the need to train classifiers on data that are sufficiently representative of heterogeneity with regards to scanning sites and mental contents.

## Acknowledgments

Data from one study (emotional memory task) were collected thanks to grants from the Canadian Institutes of Health Research, Gender and Health Institute to Dr Mendrek (CIHR grant number 200603MOP-158161-GSH-CFCA-130656). Data from 4 other studies were accessed through the SchizConnect platform (http://schizconnect.org). As such, the investigators within SchizConnect contributed to the design and implementation of SchizConnect and/or provided data but did not participate in analysis or writing of this report. Funding of the SchizConnect project was provided by NIMH cooperative agreement 1U01 MH097435. SchizConnect enabled access to the following data repository: the COllaborative Informatics and Neuroimaging Suite Data Exchange tool (COINS; http://coins.mrn.org/dx). Data from one study (resting-state) was collected at the Mind Research Network and funded by a Center of Biomedical Research Excellence (COBRE) grant 5P20RR021938/P20GM103472 from the NIH to Dr Vince Calhoun. Data from two studies (oddball and Sternberg item recognition paradigm tasks) were obtained from the Mind Clinical Imaging Consortium database. The MCIC project was supported by the Department of Energy under award number DE-FG02–08ER6458. MCIC is the result of efforts of coinvestigators from University of Iowa, University of Minnesota, University of New Mexico and Massachusetts General Hospital. Data from a fourth study (N-back task) were obtained from the Neuromorphometry by Computer Algorithm Chicago (NMorphCH) dataset (http://nunda.northwestern.edu/nunda/data/projects/NMorphCH) As such, the investigators within NMorphCH contributed to the design and implementation of NMorphCH and/or provided data but did not participate in analysis or writing of this report. The NMorphCH project was funded by NIMH grant RO1 MH056584. The last study (task-switching) was obtained through the OpenFMRI project (http://openfmri.org)from the Consortium for Neuropsychiatric Phenomics (CNP), which was supported by NIH Roadmap for Medical Research grants UL1-DE019580, RL1MH083268, RL1MH083269, RL1DA024853, RL1MH083270, RL1LM009833, PL1MH083271, and PL1NS062410. Data analysis was supported by a grant from the Natural Sciences and Engineering Research Council of Canada (NSERC grant number #436141) to PB. CD is supported by a bursary from the Lemaire foundation.

